# Multi-Omic Integration by Machine Learning (MIMaL) Reveals Protein-Metabolite Connections and New Gene Functions

**DOI:** 10.1101/2022.05.11.491527

**Authors:** Quinn Dickinson, Andreas Aufschnaiter, Martin Ott, Jesse G. Meyer

## Abstract

Cells respond to environments by regulating gene expression to exploit resources optimally. Recent advances in technologies allow the ability to gather information of cellular states of its components, measuring abundances of transcripts, their translation, the accumulation of proteins, lipids and metabolites. These highly complex datasets reflect the state of the different layers in a biological system. Multi-omics is the integration of these disparate methods and data to gain a clearer picture of the biological state. Multi-omic studies of the proteome and metabolome are becoming more common as mass spectrometry technology continues to be democratized. However, knowledge extraction through integration of these data remains challenging. Here we show that connections between omic layers can be discovered through a combination of machine learning and model interpretation. We find that model interpretation values connecting proteins to metabolites are valid experimentally and reveal also largely new connections. Further, clustering the magnitudes of protein control over all metabolites enabled prediction of gene five gene functions, each of which was validated experimentally. We accurately predicted that two uncharacterized genes in yeast modulate mitochondrial translation, *YJR120W* and *YLD157C*.We also predict and validate functions for several incompletely characterized genes, including *SDH9, ISC1*, and *FMP52*. Our work demonstrates that multi-omic analysis with machine learning (MIMaL) views multi-omic data through a new lens to reveal new insight that was not possible using existing methods.

## Main

There are various methods to integrate multi-omic datasets, reviewed in the context of single-cell data in Miao *et al*. 2021^1^. Multi-omic integration strategies currently are employed within three general disciplines: (1) disease subtyping, especially in the context of cancer heterogeneity, (2) biomarker discovery, and (3) discovery of biological insights^2^. In the context of biological insights, multi-omics integration has been accomplished using several statistical approaches, such as Bayesian, exemplified by PARADIGM^3^ and iClusterPlus^4^, or correlation-based approaches such as CNAmet^5^. These approaches have uncovered pathways involved in cancer prognosis^3^, drug selectivity of cancer lines^6^, and novel candidate oncogenes^5^.There is a need for new strategies that leverage the interactions between omics layers to produce more knowledge than the sum of the two datasets.

Machine learning is a promising approach for discovering relationships between datasets. Machine learning techniques have found success in the integration of multi-omic datasets^7^ for particular prediction tasks. Some examples of this include supervised methods predicting cancer prognosis^8^, cellular state in *E. coli*^9^, patient survival outcomes for cancer types^10^, or patient drug response^11^. Unsupervised methods have also been developed for the discovery of biomarkers^12^ and the subtyping of cancers^13^. Each of these approaches rely on an early, intermediate, or late integration strategy, as described in Picard *et al*. 2021^14^. The integration of multi-omic data through hierarchical prediction between omic layers is relatively unexplored, though at least one previous paper described prediction of metabolomic changes from proteomic changes^15^.

Here we establish multi-omic integration using a tree-based regression model trained to predict metabolite changes from proteomic changes (**Figure 1A**). This allowed us to reveal new connections between proteins and metabolites using SHAP^16^, a machine learning model interpretation method. New connections from SHAP were experimentally verified to represent the amount of control a protein’s quantity exerts over a given metabolite. Many of these protein-metabolite connections are distant based on known genetic and metabolic interactions. Finally, summarizing the strength of these protein control values across all metabolites reveals new connections between experimental conditions. In the case where conditions are single gene knockouts, this clustering reveals new functions of both characterized and uncharacterized mitochondrial proteins.

**Figure 1.**
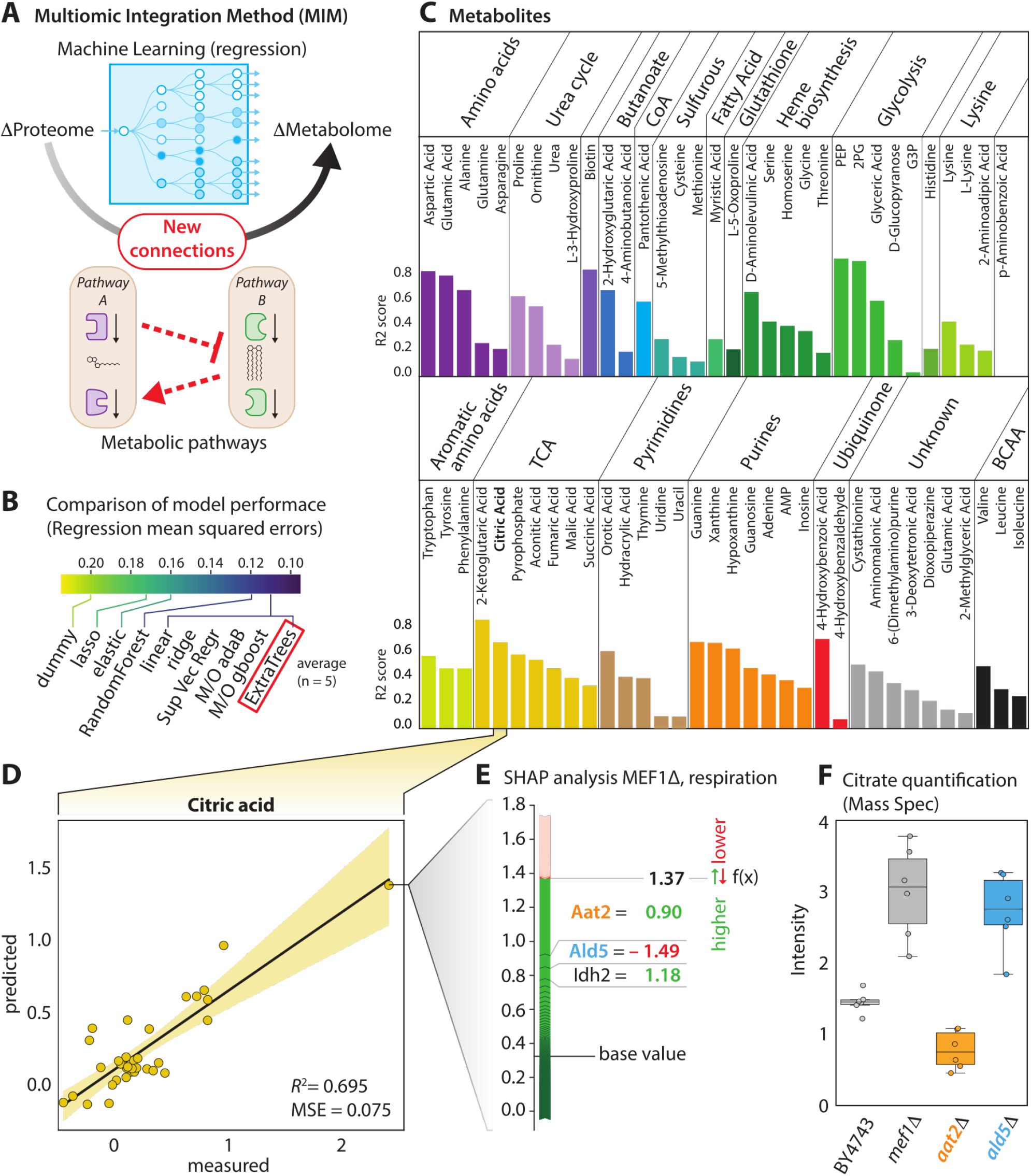
MIMaL workflow, model interpretation, and demonstration of biological applicability. **(A)** MIMAL is a multi-omic integration method utilizing machine learning model interpretation with cluster analysis to uncover unknown relationships between samples. **(B)** Comparison of the model performance with average mean squared error across 5 folds from 5-fold cross validation. ExtraTrees was selected for further analysis due to performance and specialized interpretation algorithms for decision tree based methods. **(C)** Performance of ExtraTrees models in predicting fold change in each metabolite from proteomic data, measured by R2 between predicted and experimental metabolite values for each held out test set. **(D)** Example of true versus predicted quantity of one metabolite, citric acid, with each point representing one sample, i.e. knockout strain under fermentation or respiration conditions. **(E)** SHAP forceplot for MEF1delta under respiration conditions where red and blue bars represent protein quantities that increase or decrease the prediction value of citric acid relative to the baseline, respectively. **(F)** Quantification of citric acid in strains selected from SHAP analysis. Strains were grown under respiration conditions, metabolites were extracted in methanol, and citric acid quantities were measured using targeted MS/MS. Citrate quantities reflect predictions made from SHAP.

Data was obtained from a previous multi-omic study in yeast^17^ consisting of the proteome and metabolome of wild-type or one of 174 single gene knockout yeast strains grown under fermentation and respiration conditions, for a total of 348 multi-omic profiles after computing change relative to wild-type controls. In total, the overall dataset consisted of 3,690 proteins and 273 metabolites. After imputation, data was split into training (n=313), and test (n=35) datasets. Multiple different models for each metabolite were explored (**Figure 1B**) and their performance was determined by mean squared error and R^2^ between test data model predictions and true values. The Extra Trees model was chosen as it had among the best average performance across metabolites (**Figure 1B**) and decision tree-based models have specialized model interpretation methods. Positive R^2^ scores between true and predicted quantities of metabolites in the test set were observed for nearly all identified metabolites (**Figure 1C**).

To determine the learned relationships between the proteome and metabolites, TreeSHAP was used to calculate the contribution of each protein input to the predicted level of each of the metabolites across the entire dataset. One well predicted metabolite, citric acid (R^2^ = 0.695) was chosen as an example (**Figure 1D, 1E**). The proteins with the greatest SHAP value magnitude for *mef1*Δ under respiration were Aat2 (25.46% of total magnitude) and Ald5 (4.19%) and Idh2 (3.96%) (**Figure 1F**). Unlike previous works that directly measure metabolite-protein interactions^18^, we cannot infer the nature of the interaction. We asked whether these connections reflect metabolic control by proteins by quantifying citrate in single gene knockout strains. Citrate production in *AAT2* and *ALD5* homozygous deletion mutants were compared to the BY4743 wild-type and a *MEF1* deletion mutant (**Figure 1F**) and significantly different levels of production were seen between wild type and *aat2*Δ (Student’s T-test p-value =7.22E-4), and wild type and *ald5*Δ (Student’s T-test p-value = 1.53E-3), matching the relationships predicted by the SHAP values. This result suggests that SHAP values from model interpretation may reveal protein control (ProC) over a metabolite to a greater degree than correlations (**Supplemental Figure 1**).

To further explore the relationship between proteins with the highest average ProC over citrate, GO term enrichment was performed (**Figure 2**). This analysis revealed several functional pathways that predict citrate (**Supplemental Table 1, A-D**) related to TCA cycle, stress responses, and respiration, providing further validation that these connections are biologically valid. This may also reflect the logic of the machine learning algorithm and SHAP, choosing as ProCs proteins most reflective of these functional pathways and their correlated proteins.

**Figure 2.**
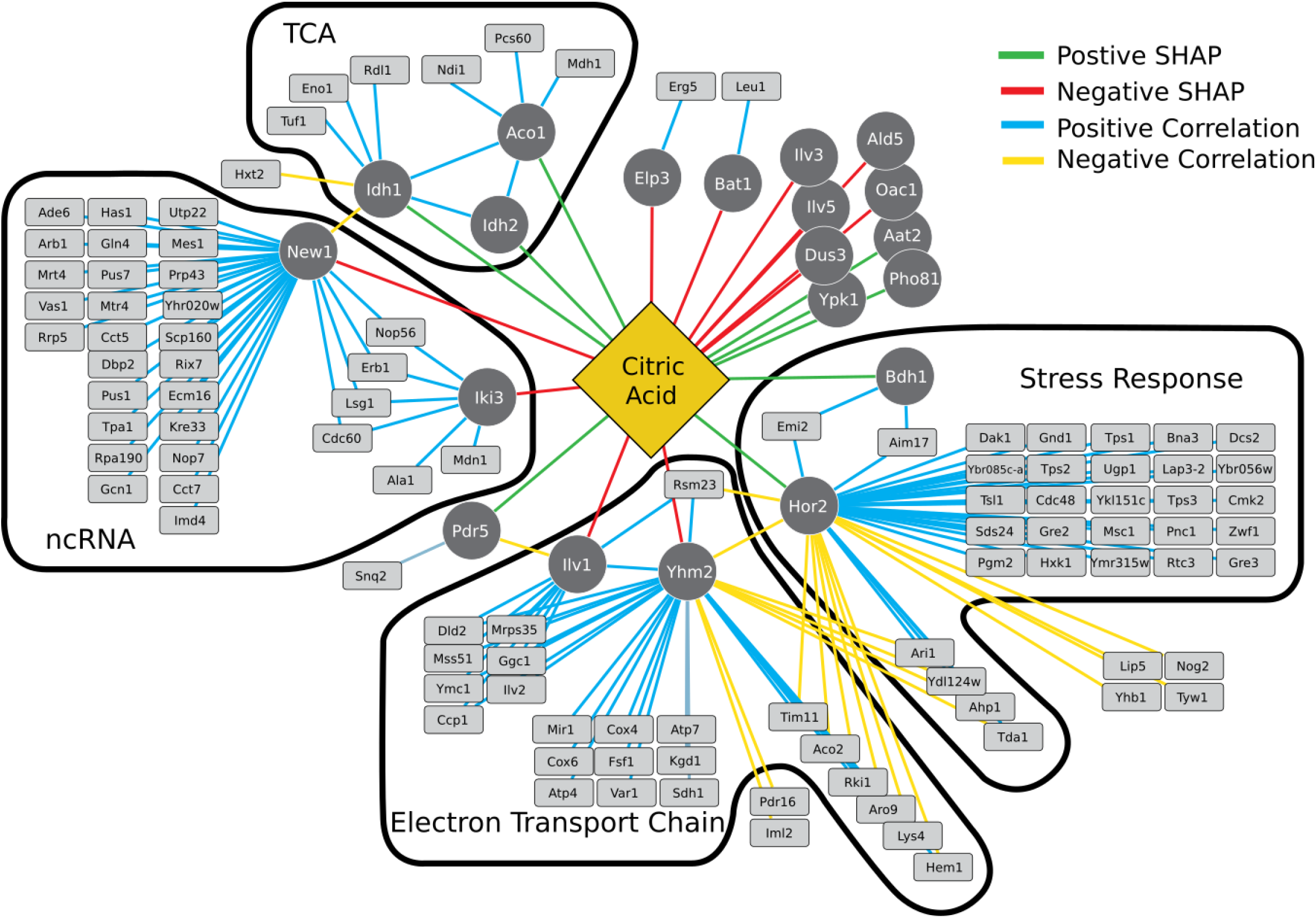
Correlated Proteins to Top 20 SHAP Values. The proteins represented by the SHAP values with the greatest average magnitude were selected for further analysis by determining their correlated proteins, to better explain the selection of these protein quantities as markers for the prediction of citric acid through the ExtraTrees model. Proteins were grouped through their shared correlations and SHAP value. Groups were examined through GO term enrichment analysis of biological processes **(Supplemental Table 1, A-D)**. Each group was labeled by a summary reflective of the other terms.

Given that our approach discovers hundreds of new connections between proteins and metabolites, we asked whether these connections are largely new or known. As expected, the top discovered connections for citrate **(Supplemental Figure 2A)** were mapped onto known positive genetic and metabolic interaction networks **(Supplemental Figure 2B)**. *AAT2, IDH1, IDH2*, and *ALD5* were close to citrate, being either one metabolic step, or one positive genetic interaction distance from an enzyme that acts directly on citrate. The remaining connections were more distant, representing new protein connections to citric acid. Notably, *OAC1, BAT1, YPK1*, and *PHO81* all lay at the median or above in calculated distance across all proteins and metabolites **(Supplemental Figure 2C)**.

Dimension reduction and clustering of ProC could reveal similarities between the input samples that were not apparent from the omic profiles alone. Because the data used here are from single gene knockouts including uncharacterized genes, this experiment tried to predict functions of the genes based on similar ProC profiles. *YDL157C* and *YJR120W* are two genes of unknown function associated with the mitochondria. Clustering of knockouts across metabolites (**Figure 3A-B**) revealed that these two knockouts frequently cluster with gene knockout strains related to mitochondrial translation (**Supplemental Table 2**). *In vivo* pulse-chase radiolabeling of mitochondrial translation in wild type and *ydl157c*Δ and *yjr120w*Δ revealed changes in mitochondrial translation (**Figure 3C, Supplemental Figure 3, A-B**). *ydl157c*Δ had a global reduction of mitochondrial translation and absence of *YJR120W* resulted in a dysregulation of translation. In *yjr120w*Δ, Var1, Cox2, Cox3, and Atp6 are down regulated, with more pronounced downregulation seen for Cox3 and Atp6. Cyt *b* however was upregulated. This alteration in translation might reflect previously suggested interactions of *YJR120W. YJR120W* is upstream of *ATP2* on the yeast chromosome, and the deletion of *YJR120W* was previously noted to alter *ATP2*’s expression.^17^ Atp2 is a part of the F_1_sector of the F_1_F_o_ATP synthase, which regulates the mitochondrial translation of *ATP6* and *ATP8*^19^. In line with these observations, deletion of *YDL157C* severely impaired respiratory growth, while the effect of the deletion of *YJR120W* was less pronounced (**Supplemental Figure 3C**).

**Figure 3.**
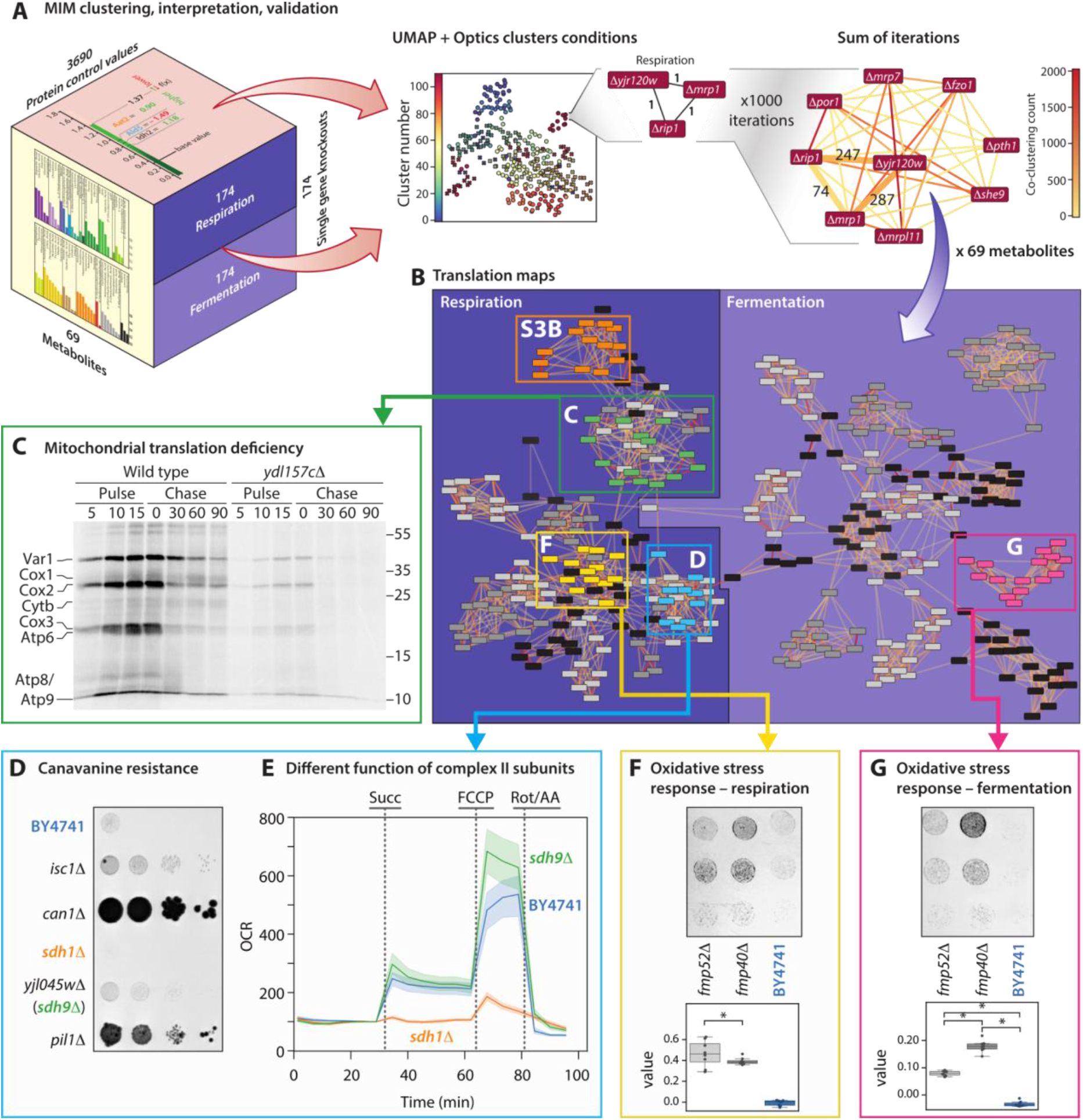
MIMaL clustering, interpretation, and validation. **(A)** Overview of the method to find connections between conditions using dimensionality reduction, clustering, and network analysis. SHAP values were calculated for all proteins across all knockouts. UMAP was used to reduce dimensionality to 10 dimensions, the first two are displayed graphically. UMAP dimensions were clustered with OPTICS. UMAP and OPTICS were repeated 1000 times for each metabolite. **(B)** A graph was constructed where each edge is linearly proportional to the count of co-clustering across the 69,000 clustering repetitions and with a minimum cutoff for including edges. **(C)** Autoradiographic image of gel assessing mitochondrial translation in wild type and *ydl157cΔ* cells were treated with cycloheximide and using 35S-methionine for 15 minutes (pulse) at 30°C. The labeling was stopped by adding excess cold methionine and the temperature was increased to 37°C to induce protein destabilization (chase for a total of 90 min). **(D)** Resistance to canavanine stress. Strains were grown on synthetic complete media minus arg +2.5 µg/ml canavanine for 18 days. *ISC1* and *SDH9* knockout strains were connected to *pil1Δ*, which was previously shown to resist canavanine stress like *can1Δ* (positive control). Both *isc1Δ* and *sdh9Δ* showed resistance to canavanine compared to wild type. **(E)** Oxygen consumption in responses to succinate as the sole carbon source measured by seahorse respirometry. Responses to succinate were significantly different (p -value = 0.001, Tukey’s HSD) between *SDH1* and *SDH9* knockouts. **(F)** The strains *fmp40Δ* and *fmp52Δ* were tested for resistance to hydrogen peroxide stress under respiration **(F)** and fermentation **(G)** conditions and compared using image analysis of drop dilution assays **(Supplemental Figure 3G)**. Differences between all strains was significant (p-value = 0.001, Tukey’s HSD) under fermentation conditions, and significant (p-value = 0.001, Tukey’s HSD) between wild-type and the others under respiration conditions. Drop dilution image colors are inverted to enhance visibility.

Although the connections between translation and *YDL157C* and *YJR120W* were not discovered in the original paper that reported this data, closer inspection of correlation between proteome profiles resulting from gene knockouts may have revealed this relationship. We wondered if our summary strategy of ProC values could reveal new gene connections that would not be apparent from an omic profile similarity alone. To further test the relationships predicted by the clustering network, three additional clusters were analyzed for their connections to incompletely characterized genes. The first of these clusters included *YJL045W*, now annotated as *SDH9* as it is a paralog of *SDH1*. Unexpectedly, *sdh9*Δ was found to lack direct connections to *sdh1*Δ under respiration conditions in the final trimmed network but rather had the greatest connection to Pil1, a key protein in eisosomal structure^20^. The eisosome is a membrane structure involved in membrane transport. One transporter associated with the eisosome is Can1, an arginine transporter whose deletion confers resistance to the toxic, non-proteinogenic amino acid canavanine^21^. Disruption of the eisosome through deletion of *PIL1* has also been shown to provide resistance to canavanine^22^. To test the connection between *SDH9* and the eisosome, the growth of deletion strains of *SDH9, SDH1, CAN1, PIL1*, and another connection to *PIL1, ISC1*, were tested on synthetic complete media (SC) without arginine + canavanine. All tested strains, other than *sdh1*Δ, which had a growth defect on SC -arg (**Supplemental Figure 4A**), were shown to grow in the presence of canavanine better than wild type (**Figure 3D**). Additionally, all strains but *pil1*Δ showed significantly higher viability when exposed to very high concentrations of canavanine over 72 hours (**Supplemental Figure 4B**). However, as *sdh1*Δ showed a growth defect on SC -arg, the link between *SDH1* and eisosomal function remains ambiguous.

We wondered why SDH9, a gene annotated to function in complex II, would convey resistance to canavanine. To test the link between SDH1 and SDH9, respiratory responses were quantified; we used succinate as a source of electrons to complex II and SDH9Δ showed a response more similar to wild type than SDH1Δ. Oxygen consumption rate (OCR) spiked in SDH9Δ when exposed to succinate, while this was not observed in SDH1Δ. (**Figure 3E**). The different responses to succinate demonstrate the distinctiveness of the two succinate dehydrogenases and suggest unique functions for each.

Also of note is the resistance of *isc1*Δ to canavanine. Isc1 is an enzyme involved in sphingolipid hydrolysis to ceramides^23^ and is activated by cardiolipin^24^. Proteins involved in cardiolipin biosynthesis are significantly enriched in the cluster containing *isc1*Δ and *pil1*Δ (**Supplemental Table 1E**). This supports an interplay between cardiolipin, ceramides, and the eisosome as suggested in the literature^24,25^, which should be explored further in future studies.

The final two clusters analyzed include another uncharacterized gene in both respiration and fermentation conditions, *FMP52. fmp52*Δ was found to have the greatest connection weight to *fmp40*Δ. Fmp40 is an AMPylator involved in the oxidative stress response^26^. In addition, Fmp52 had the second greatest connection weight to Aim25, a protein of unknown function involved in the oxidative stress response^27^. Based on these connections, it seemed likely that *fmp52*Δ would have an altered response to oxidative stress and therefore show a difference in resistance to oxidative stressors, such as hydrogen peroxide. To test this hypothesis, cells under respiration and fermentation conditions were exposed to hydrogen peroxide and their viability was determined after 30 minutes (**Figure 3F, 3G, Supplemental Figure 5A**). The resistance to hydrogen peroxide was significantly higher in both *FMP40* and *FMP52* deletion strains compared to WT controls. Under fermentation conditions, there was a significant difference between the resistance of *fmp40*Δ and *fmp52*Δ, while under respiration conditions there was no significant difference. This coincides with the weight of the connections between *FMP40* and *FMP52* in the network; the weight of the edge connecting them is substantially larger in the respiration cluster. As a separate test, *fmp40*Δ and *fmp52*Δ were grown under respiration conditions in a zone-of-inhibition assay with hydrogen peroxide. A similar result was found, with both the *fmp40*Δ and *fmp52*Δ lawns growing closer to the source of hydrogen peroxide (**Supplemental Figure 5B**).

To compare the performance of this clustering method with proteomic correlations, we looked at the representation of known genetic and physical interactions among the top selected connections (**Supplemental Table 3**) from the clustering analysis and the correlations between proteomes of knockout strains. As an example, of the 873 known genetic and physical interactions between the genes represented by the knockout strains under fermentation conditions, 45 were uniquely represented across all proteomic correlations, 31 shared by correlations and clustering, and 85 uniquely represented by clustering analysis (**Supplemental Figure 6**).

In conclusion, we have shown that SHAP model explanation values can reflect true biological relationships between the proteome and metabolome, demonstrated the novel application of SHAP model explanation values in the integration of multi-omic data, and illustrated the utility of this method through the characterization of several uncharacterized yeast genes. We foresee this method being useful as a new multi-omic integration technique that provides unique insight into the relationships between multi-omic levels. Additionally, the data we have generated may act as a reference to better understand orphan mitochondrial proteins.

## ONLINE METHODS

### Yeast protein-metabolite machine learning

A total of 873 proteins were measured in all samples. Missing protein values were imputed using the sklearn function KNNImputer with setting n_neighbors=2, resulting in all 3,690 protein quantities being used as input for the modeling task. Metabolite data was imputed using the same setting, producing 273 complete metabolite columns.

### Machine Learning and Model optimization

The data was split into 313 random examples for training and 35 examples for testing. Multiple types of models were first tested 5-fold cross validation with the default parameters, and the average mean squared error (MSE) across the five folds was compared (see jupyter notebook on github ‘compare-models.ipynb’). Tested models were implemented in sklearn including: a dummyRegressor baseline, LinearRegression, Lasso, ElasticNet, Ridge, Support vector regression wrapped in MultiOutputRegressor, AdaBoost^28^ wrapped in multiOutputRegression with 500 estimators, GradientBoostingRegressor with 500 estimators wrapped in MultiOutputRegressor, ExtraTreesRegressor with 500 estimators, and RandomForestRegressor with 500 estimators^29^. All of these models except the dummy, elasticNet, and Lasso performed similarly according to the metric MSE; we selected extraTreesRegression because we wanted the interpretability of a tree model and the speed of training extraTrees.

One multi-output regression extra trees model was optimized using 5-fold cross validation with the 313 training examples by gridsearch (see jupyter notebook on github “extratrees-gridsearch.ipynb”) with the following parameters: ‘max_depth’: [10,30, 50, 70,None], ‘min_samples_leaf’: [1,2,5], ‘min_samples_split’:[2,5,10], ‘max_features’: [‘log2’, ‘auto’, ‘sqrt’], ‘n_estimators’: [500, 1000, 1500]. The best model parameters for the polar metabolomics model used all the default parameters except max_depth=50 and n_estimators=500. Those parameters were then used to train a single output extraTrees model for each of the 273 polar metabolites. The trained model was used to make predictions on the 35 examples in the test set, and those true and predicted values were used to compute regression metrics. The R2_score and mean_square_error functions in sklearn summarized performance across all the metabolites.

### Yeast Protein-Metabolite SHAP Analysis

SHAP values were calculated for each knockout for each metabolite model using TreeExplainer. Only identified metabolites that had a positive R^2^ score comparing the true versus predicted quantity were included in subsequent analysis. This excludes roughly 200 additional unidentified metabolites.

Correlations between each protein quantity across all single knockout samples was calculated using Spearman’s rho and significance was adjusted using Bonferroni Correction. For citric acid, the top 20 mean magnitude SHAP contributor proteins were chosen for further analysis. A network was created with citric acid as the central node, linked to each SHAP contributor protein. Each SHAP contributor protein was then linked to each correlated protein with an adjusted p-value < 0.05 and a |ρ|>0.7. Finally, correlations between correlated proteins were labeled with edges. Enrichment analysis was performed using ClueGO on each group of SHAP contributor proteins sharing positive correlations and their positively correlated proteins.

### Citrate quantification by direct infusion MS/MS

Yeast strains were grown overnight in YPD at 30°C. After growth, OD595 was measured and cells were washed with PBS. YPDG was inoculated to an initial OD595 of 0.01 and grown at 30C for 24 hours. After growth, OD595 was measured and the equivalent of 0.37 OD595 at 1ml was harvested from each. These cells were pelleted, washed with PBS, pelleted, frozen with LN2, and stored at -80C. To extract metabolites, each pellet was resuspended in 185 ul 75% methanol, placed at 100C for 5 minutes, vortexed for 30 seconds, and cooled on ice. Cell debris was pelleted and supernatant was used for citrate quantification.

Mass spectrometry was performed on a Thermo Scientific Exploris 240, using a Thermo Scientific Nanospray Ion Source. One ul of each extract was directly infused into the mass spectrometer. To quantify citrate, targeted MS/MS was performed, targeting the ion at 191.0192 m/z. The measured intensity of the fragment at 111.008 m/z was integrated across 811 scans to determine total citrate present in each sample. Data analysis was performed using pyteomics.

### Clustering Metabolite Control of Knockout Yeast to Predict Gene Function

SHAP values of the knockouts were clustered using a combination of Uniform Manifold Approximation and Projection (UMAP)^30^ and Ordering Points To Identify Cluster Structure (OPTICS)^31^ to determine clustering and likely function of unknown mitochondrial genes. For UMAP, dimensionality of data (n_components) was set at 10, neighbors (n_neighbors) was set to 3, minimum distance (min_dist) was set to 0, and the distance metric (metric) was manhattan. For OPTICS, minimum samples (min_samples) was set to 2. All other parameters were set to their defaults.

To generate the final clusters and account for the stochasticity of UMAP, UMAP and OPTICS clustering was repeated 1000 times for each metabolite. The clusters generated from each repetition were compared by creating a network with each node representing one of the knockouts and each weighted edge representing twice the number of times the knockouts clustered together of the 1000 repetitions.

The weighted edges, representing the membership of clusters, were combined across known, non-repeated metabolites with a model performance of R^2^>0. To determine a subset of most relevant connections, a linear regression was calculated between the edge weight and the rank of the edge when sorted in descending order. All edges with a weight that lay above the linear regression (a weight of 8210) were included as the relevant connections. Nodes were clustered in Cytoscape using the Markov Cluster Algorithm (MCL Cluster in clusterMaker). Layout of the network was calculated using the Prefuse Force Directed Layout.

### Known Connections network

To create the yeast metabolic network, a list of reactions, enzymes, compounds, and enzymatic reactions was downloaded from Biocyc^32^. These datasets were combined to create a metabolic network consisting of all known pathways and their associated enzymes. The following nodes and associated edges were removed from the network due to their ambiguity and relative abundance across reactions: “PROTON”, “WATER”, “ATP”, “ADP”, “PPI”, “Pi”, “Protein-L-serine-or-L-threonine”, “Protein-Ser-or-Thr-phosphate”, “AMP”, “NAD”, “NADH”, “CO-A”, “NADP”, “NADPH”, “CARBON-DIOXIDE”, “GLT”, “S-ADENOSYLMETHIONINE”, “OXYGEN-MOLECULE”, “ACETYL-COA”, “AMMONIUM”, “ADENOSYL-HOMO-CYS”, “Nucleoside-Triphosphates”, “Peptides-holder”, “RNA-Holder”, “Cytochromes-C-Oxidized”, “Cytochromes-C-Reduced”, “GDP”, “Ubiquitin-C-Terminal-Glycine”, “General-Protein-Substrates”. Edges between enzymes and compounds were assigned a weight of 3.

A list of all known *S. cerevisiae* positive genetic interactions was downloaded from Saccharomyces Genome Database (SGD). Every ORF absent from the network, i.e. those whose protein does not catalyze a metabolic reaction, were added as nodes and edges with a weight of 10 were created to link ORF nodes with known positive interactions. Weighted closest distance to citrate was calculated for every node using Dijkstra’s algorithm. The closest distance can be summarized as 3 + 6*(metabolic distance) + 10*(positive interaction distance)

### Comparison of Clustering to Proteome Profile Correlation

A list of all possible pairwise combinations of the 174 proteins represented by the knockout strains was generated. A set of all known genetic and physical interactions for the 174 genes was downloaded from the Saccharomyces Genome Database. For each pairwise combination, it was determined if the pair was correlated through proteomic data, connected through clustering analysis, and if it had known genetic or physical interactions. The overlap of correlations and clustering connections with known interactions was determined and plotted using matplotlib-venn.

### Yeast strains and genetics

All strains used for translation assays were isogenic to *Saccharomyces cerevisiae* W303 *MAT* a {*leu2-3,112 trp1-1 can1-100 ura3-1 ade2-1 his3-11,15*} obtained from Euroscarf and are listed in **Supplementary Table 4**. Chromosomal modifications were made by PCR-based amplification of cassettes followed by integration via homologous recombination, according to Janke et al^33^ and applying lithium acetate transformation according to Gietz and Woods^34^. All plasmids and oligonucleotides used for this approach are listed in **Supplementary Table 5**. Transformants were validated via growth on selection media and PCR-based confirmation of locus-specific integration.

Strains for the other assays were in BY4743 background for the citrate quantification or BY4741 for the canavanine and hydrogen peroxide assays. All strains were obtained from Horizon Discovery and are listed in **Supplementary Table 4**.

### Translation Assay - Media and culturing conditions

Strains were cultivated at 30°C and 170 rpm shaking. Full media (YEP) contained 1% yeast extract (Bacto, BD Biosciences), 2% peptone (Bacto, BD Biosciences) and 2% glucose, 2 % galactose or 2% glycerol as carbon source. Synthetic complete (SC) media consisted of 0.17% yeast nitrogen base (Difco, BD Bioscience). 0.5% (NH_4_)_2_SO_4,_ 20 mg/l adenine, 20 mg/l uracil, 20 mg/l arginine, 15 mg/l histidine, 30 mg/l leucine, 30 mg/l lysine, 15 mg/l tryptophan, 30 mg/l isoleucine, 20 mg/l methionine, 50 mg/l phenylalanine, 20mg/l threonine, 20 mg/l tyrosine, 150 mg/l valine and carbon sources as indicated above. All components were separately prepared in distilled water, autoclaved (25 min, 121°C, 210 kPa, except histidine and tryptophan, which were sterile filtered using 0.2 µm filters) and mixed before use. For solid media, 2% agar was admixed.

### *In vivo* labeling of mitochondrial translation products

[^35^S]-methionine-based *in vivo* labeling of mitochondrial translation products was performed according to Carlström et al^35^ with slight modifications. Cells were grown in SC medium containing galactose as carbon source (SC-Gal) to mid-logarithmic phase (approx. OD^600^ = 1.5 -2) and washed three times in 5 ml H_2_O. Strains were subsequently washed once in 5 ml SC-Gal media without amino acids and a volume corresponding to OD_600_ = 4 was harvested and resuspended in 1.5 ml SC-Gal media without amino acids. Amino acids were admixed (18 µg of each amino acid, without methionine) and incubated for 10 min at 30°C, 600 rpm shaking. To stop cytosolic translation, cycloheximide was added to a final concentration of 150 µg/mL and incubated for 2.5 min at 30°C, 600 rpm shaking. 3 µl of [^35^S]-methionine (10 mCi/ml) were added to start the labeling reaction. For pulse-labeling, 200 µl aliquots were harvested after 5, 10 and 15 min, mixed with 50 µl of Stop solution (1.85 M NaOH; 1 M β-mercaptoethanol; 20 mM PMSF) and 10 µl of 200 mM cold methionine, and placed on ice. To follow stability of newly synthesized mitochondrial proteins, 40 µl of 200 mM cold methionine was added to the remaining cell suspensions and incubated at 37°C, 600 rpm (chase). Thereby, 200 µl samples were harvested 30, 60 and 90 min after addition of cold methionine, mixed with stop solution as described above and placed on ice.

### SDS-PAGE and Immunoblotting

Trichloroacetic acid was added to [^35^S]-methionine-labeled samples with a final concentration of 14%, incubated for 30 min on ice and subsequently centrifuged for 30 min, 20.000 g at 4°C. Supernatants were carefully removed and pellets rinsed once in 1 ml 100% acetone. After further centrifugation for 30 min at 20.000 g at 4°C, supernatants were removed and pellets resuspended in 75 µl sample buffer (50 mM Tris-HCl, 2% SDS, 10% glycerol, 0.1% bromophenol blue, 100 mM DTT; adjusted to pH 6.8). Subsequently, samples were incubated for 10 min at 65°C, 1400 rpm shaking. 30 µl of the sample were loaded on 16%/0.2% SDS polyacrylamide/bis-acrylamide gels. After separation, proteins were transferred to a nitrocellulose membrane, which was stained with Ponceau S. Protein standard bands (PageRuler^TM^ Plus Prestained Protein Ladder, ThermoFisher) on the nitrocellulose membrane were marked with diluted [^35^S]-methionine solution and the membranes were applied for autoradiography. Detection was performed with a Fujifilm FLA-9000 phosphorimager.

Membranes were subsequently applied for immunoblotting, using Mrp1^36^, Mrpl36^37^ and Tom70 (kind gift from Prof. Rapaport, University of Tübingen) specific antibodies, as well as anti-rabbit secondary antibody (Sigma, A0545).

### Drop dilution assay

To monitor cellular growth, yeast strains were cultivated in YEP media containing either glucose or glycerol to mid-logarithmic phase (approx. OD_600_ 1.5-2). Cultures were washed three times in YEP media without carbon source and a volume corresponding to OD_600_ = 1 was harvested. Samples were resuspended in 1 ml YEP media without carbon source and three serial 1:10 dilutions thereof were created. 3 µl of cell suspensions were spotted on YEP agar plates either containing glucose or glycerol as carbon source. Plates were incubated for 2 days at 30°C and photographed with a VWR GenoPlex system.

### Canavanine Drop Dilution

Cultures were grown for 18 hours in 1ml YPD for BY4741 or YPD + G418 for the knockout strains. Cultures were centrifuged at 3000 rcf for 3 minutes and pellets were resuspended in 3ml YPG. After 24 hours, the cultures were pelleted, washed with SC -Arg +glycerol and adjusted with SC -Arg +glycerol to an OD660 of 0.1 and plated onto SC - Arg +glycerol or SC - Arg +glycerol +canavanine at 0.25 μg/ml plates with dilutions of 1, 1:10, 1:100, 1:1000, 1:2000, 1:4000, 1:8000, and 1:16,000. Plates were incubated at 30°C and pictures were taken after 1 week and again at 18 days. Images of colony formation were captured using ImageLab software with a Bio-Rad GelDoc.

### Canavanine Viability

Cultures were grown for 18 hours in 1ml YPD or YPD + G418. Cultures were centrifuged at 3000 rcf for 3 minutes and pellets were resuspended in 3ml YPG. After 24 hours, the cultures were pelleted, washed with SC -Arg +glycerol and adjusted with SC -Arg +glycerol to an OD660 of 0.2. 100 μl was adjusted with SC -Arg +glycerol to an OD660 of 0.1 and plated onto YPD plates with dilutions of 1, 1:10, and 1:100 and refrigerated at 3°C for 72 hours. The remaining culture was adjusted to an OD660 of 0.1 with SC -Arg +glycerol +1200 μg/ml canavanine(final concentration 600 μg/ml) and incubated with shaking at 30°C for 72 hours. Cultures OD660 were centrifuged, washed, and adjusted to 0.1 OD with SC -Arg +glycerol. Cultures were then plated onto the previously refrigerated YPD plates at dilutions of 1, 1:10, and 1:100. Plates were incubated at 30°C for 18 hours. Images of colony formation were captured using ImageLab software with a Bio -Rad GelDoc.

### Hydrogen Peroxide Viability

Cultures were grown for 18 hours in 2ml YPD or YPD + G418. 1ml of each culture was centrifuged at 3000 rcf for 3 minutes and pellets were resuspended in 3ml YPG and incubated for 24 hours at 30°C. To the remaining preculture, 2ml YPD was added and incubated at 30°C for 5 hours. For each set of cultures after incubation, the cultures were pelleted, washed with YPD or YPG and adjusted with YPD or YPG to an OD660 of 0.2. For fermentation, 100 μl of each culture was added to 100µl YPD or YPD + 128 mM hydrogen peroxide. For respiration, 100µl of each culture was added to 100µl YPG or YPG +1024 mM hydrogen peroxide. Cultures were exposed to hydrogen peroxide for 30 minutes. After treatment, cells were plated onto YPD plates at dilutions of 1, 1:10, 1:100, 1:1000. Plates were incubated for 18h at 30°C. Images of colony formation were captured using ImageLab software using a bio-rad GelDoc.

### Quantification of Images

To quantify the growth of the drop dilution assays, images were exported in the TIF format at a DPI of 600. ImageJ was used to measure the brightness (R+G+B)/3 of circles 0.015 in^2. Six circles were used as a background and circles measuring the drops were centered. Circles were drawn after each measurement to mark each location (**Supplemental Data**). To calculate growth ratios, the average background measurement was subtracted from each brightness measurement. Then the experimental brightness was divided by the control brightness for each strain to calculate a ratio of growth. Average ratios were plotted in seaborn and differences between strains were compared using ANOVA and Tukey’s Post Hoc test.

### Hydrogen Peroxide Zone of Inhibition

Cultures were grown for 18 hours in 1ml YPD or YPD + G418. 1ml of each culture was centrifuged at 3000 rcf for 3 minutes and pellets were resuspended in 3ml YPG and incubated for 24 hours at 30°C. The OD660 was adjusted to 1 for each culture. 1ml of culture was plated onto 25ml YPG plates and allowed to dry. To create the hydrogen peroxide gradient, a central section of each plate was removed using a 1ml pipette tip. 100µl 3% hydrogen peroxide was added to the central hole and allowed to diffuse. Plates were incubated for 1 week at 30□C. Images of lawn formation were captured using ImageLab software using a bio-rad GelDoc.

### Seahorse Assay

To prepare the seahorse plate, 50 ul of poly-L-lysine (0.1 mg/ml) was added to each well and allowed to sit for 2 hours. The solution was aspirated and washed with 100ul sterile water. The coated plate was stored at 3C until ready for the assay. On the day of the assay, the plate was brought to room temperature and 80 ul of seahorse media was added to each well. An additional 100 ul of seahorse media were added to wells acting as baselines. Injections were prepared to have a final concentration of 5mM ethanol or succinate, 1uM FCCP, 1uM rotenone, and 1uM antimycin A.

To prepare cells for the seahorse assay, cells were grown overnight in 1 mL YPD. After growth to stationary phase, cells were pelleted and resuspended in 4 mL YPG. Cells were grown for 25 hours. Cells were pelleted and resuspended in seahorse media (6.6g/l YNB + NH4SO4) to a final OD660 of 0.38. Each sample was diluted an additional 1:5 in seahorse media and 100 ul of culture were placed into each well of the prepared seahorse plate. The plate was centrifuged at 250 rfc for 3 minutes and incubated at 30C for 30 minutes.

Plates were measured on a Seahorse XF-96. A total of 18 measurements of the oxygen consumption rate (OCR) and extracellular acidification rate (ECAR) were taken over 96 minutes, with 10 technical replicates for each strain. Six initial measurements were taken as a baseline, six measurements were taken after the injection of succinate, three measurements after the injection of FCCP, and three final measurements after the injection of rotenone/antimycin A. Data collected was analyzed using Agilent Wave and pandas and plotted using seaborn and matplotlib.

## Code and data availability

The code and data are available at https://github.com/jessegmeyerlab/MIMaL.

Supporting data is available from zenodo (https://doi.org/10.5281/zenodo.6537297).

### Supporting data

https://doi.org/10.5281/zenodo.6537297

### Supporting Information 1. H_2_O_2_ Curve

This folder contains images of yeast colonies from left to right, BY4741, *fmp40Δ, fmp52Δ*, and *set4Δ* after treatment with several concentrations of hydrogen peroxide under both respiration and fermentation conditions.

### Supporting Information 2. Canavanine Viability

This folder contains images of yeast drop dilution colonies of strains, from left to right, BY4741, *isc1Δ, can1Δ, sdh1Δ, sdh9Δ*, and *pil1Δ* on YPD plates after 24h. Rows 1, 3, and 5 were treated for 72h with canavanine. Black circles mark areas used for quantification by measuring grayscale for **Supplemental Figure 4 C**. Original pictures are in the folder originals.

### Supporting Information 3. Canavanine Drop Dilution

This folder contains images of yeast drop dilution colonies of strains, from left to right, BY4741, *isc1Δ, can1Δ, sdh1Δ, sdh9Δ*, and *pil1Δ* on SC -arg plates with or without 2.5ug/ml canavanine after one week or 18 days. Rows represent 1:10 dilutions from top to bottom.

### Supporting Information 4. H_2_O_2_ Zone of Inhibition

This folder contains images of yeast lawns from each strain, BY4741 (1, 5, 9, 13), *fmp40Δ* (2, 6, 10, 14), *fmp52Δ* (3, 7, 11, 15), and *set4Δ* (4, 8, 12, 16) after growth on YPG with hydrogen peroxide added to the center of each plate.

### Supporting Information 5. H_2_O_2_ Viability

This folder contains images of yeast drop dilution colonies of strains, from left to right, BY4741, *fmp40Δ, fmp52Δ*, and *set4Δ* and repeated on YPD (fermentation folder) or YPG (respiration folder) plates after 24h. The right half were treated for 30 minutes with hydrogen peroxide. Black circles mark areas used for quantification by measuring grayscale for **Figure 3 F**,**G**. Original pictures are in the folder originals.

### Supporting Information 6. Cytoscape Network

This file is the full network for **Supplemental Figure 2 C**. This includes labels and distances for all nodes.

## Supplemental Tables

### Supplemental Table 1. GO enrichment

These tables consist of significantly enriched GO terms for each group of correlated proteins listed in **Figure 2** and of the cluster containing PIL1Δ_resp in **Figure 3 B**. For each table, the GO ID, Term, Ontology Source, term p-value, term corrected p-value, group p-value, group corrected p-value, GO level, GO group, percentage of associated genes, number of associated genes, and the associated gene names are listed. The first tab contains the GO terms associated with the ncRNA group. The second tab contains the GO terms associated with the Electron Transport Chain group. The third tab contains the GO terms associated with the Stress Response group. The fourth tab contains the GO terms associated with the TCA Cycle group. The fifth tab contains the GO terms associated with the *pil1*Δ_resp cluster.

### Supplemental Table 2. Network Clusters

This table contains the membership of each condition, i.e. knockout strain under fermentation or respiration, to each cluster present in **Figure 3 B**.

### Supplemental Table 3. Selected Connections

This table contains the list of all connections and associated weights selected through the linear regression method (**Supplemental Figure 4A**) and displayed in **Figure 3 B**.

### Supplemental Table 4. Yeast Strains Used in this Study

This table consists of the names of all *S. cerevisiae* strains used in this study, along with their genotype, and their source.

### Supplemental Table 5. Oligonucleotides and PCR templates used for chromosomal modification and respective control experiments

This table lists the modification intended by each listed oligonucleotide, the oligonucleotide’s sequence, and the PCR source for each oligonucleotide.

The authors declare no conflicts of interest.

#### ACKNOWLEDGEMENTS

This work was supported by the United States National Institute of Health (NIH) NIGMS R35 GM142502, the National Institute on Aging (NIA) R21 AG074234, the Swedish research council and the Knut and Alice Wallenberg foundation. This research was completed in part with computational resources and technical support provided by the Research Computing Center at the Medical College of Wisconsin. We thank Monika Zielonka and the Redox & Bioenergetics Shared Resource at the Medical College of Wisconsin Cancer Center for help with seahorse data collection. We thank H. Adam Steinberg for graphic design help. We thank Jong -In Park for helpful discussions. We thank Yuming Jiang for help with experiments.

## CONTRIBUTIONS

Conceptualization QD, JGM

Methodology QD, JGM

Software QD, JGM

Validation QD, JGM

Formal Analysis QD, JGM, AA

Investigation QD, JGM, AA

Resources, JGM, MO

Data Curation QD, JGM

Writing - Original Draft QD, JGM

Writing - Review and Editing QD, JGM, MO

Visualization QD, JGM

Supervision JGM, MO

Project Administration JGM

Funding Acquisition JGM, MO

---

**Supplemental Figure 1.**
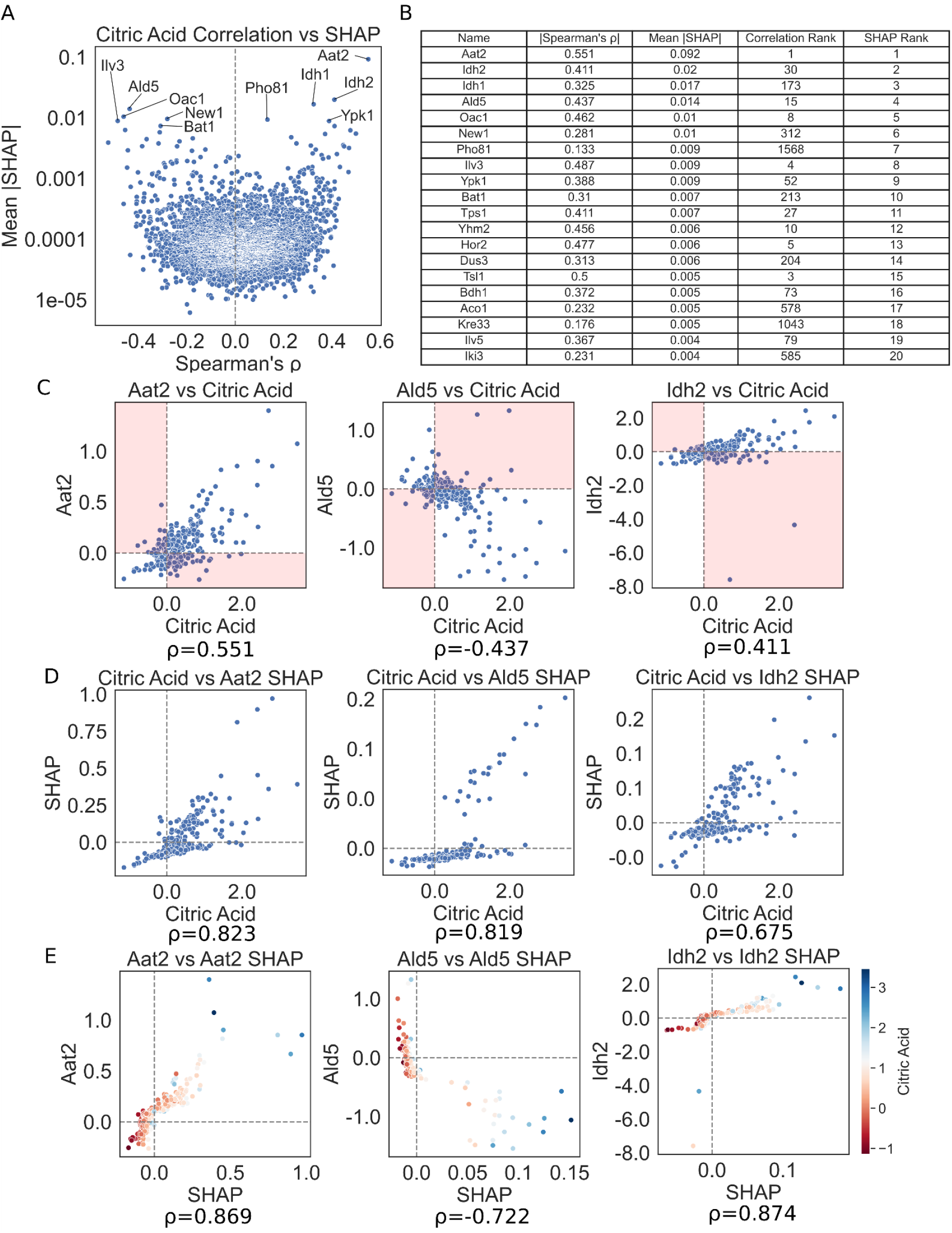
Correlations and SHAP Values as Measures of ProC Over Citric Acid. **(A)** Mean absolute SHAP values for each protein are plotted with each protein’s Spearman correlation with citric acid. The top ten magnitude SHAP proteins are labeled. **(B)** Ranking of proteins by mean absolute SHAP values and absolute Spearman correlations are compared. Although Aat2 is both the top rank for SHAP and correlation, the other top ranked SHAP proteins lay considerably lower, illustrating that the discovery of ProC of metabolites through SHAP discovers relationships beyond simple correlations. **(C)** Correlation of citric acid and protein values with members of the top ProC for citric acid. The usage of a linear correlation between proteins and a metabolite as a ProC can be inaccurate due to examples where the correlation is false, highlighted in red.**(D)** Correlation of SHAP values with citrate levels. Higher correlations are seen between SHAP and citrate than looking at direct protein folds. **(E)** Correlation of protein values with SHAP contributions of members of the top ProC for citric acid. While correlated, the SHAP values show a non-linear relationship with protein concentrations at the outliers, adjusting the relationship to better predict citrate.

**Supplemental Figure 2.**
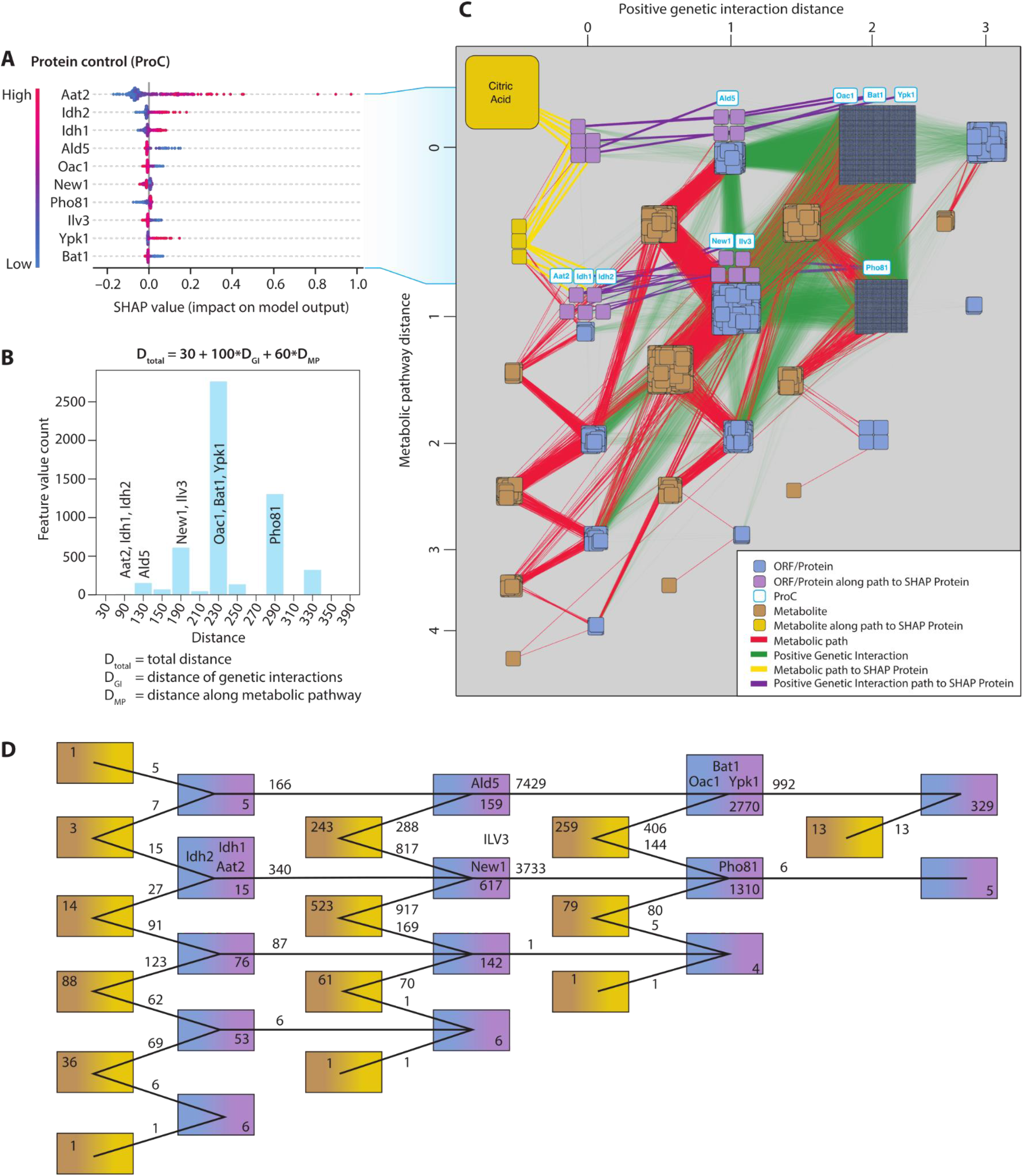
MIMaL Reveals New Connections Between Proteins and Metabolites. **(A)** Top 10 SHAP values for citric acid across all conditions, sorted by mean magnitude SHAP by each protein. **(B)** A network consisting of the metabolic pathways present in S. cerevisiae was constructed from data obtained from Biocyc. Connections between proteins through metabolites have a weight of 6. Positive genetic interactions among all ORFs in yeast were downloaded from Saccharomyces Genome Database and added to the network with a weight of 10. Distance to citrate was calculated using the Dijkstra algorithm, and can be represented by 3+(# genetic interactions)*10 + (# metabolic reactions)*6. **(C)** The overall distribution of distances were plotted as a histogram. The network was organized by distance to citrate and the proteins representing the top 10 SHAP values for citric acid prediction were highlighted, along with their paths to citric acid. **(D)** A representation of the total number of nodes and connections between each category.

**Supplemental Figure S3.**
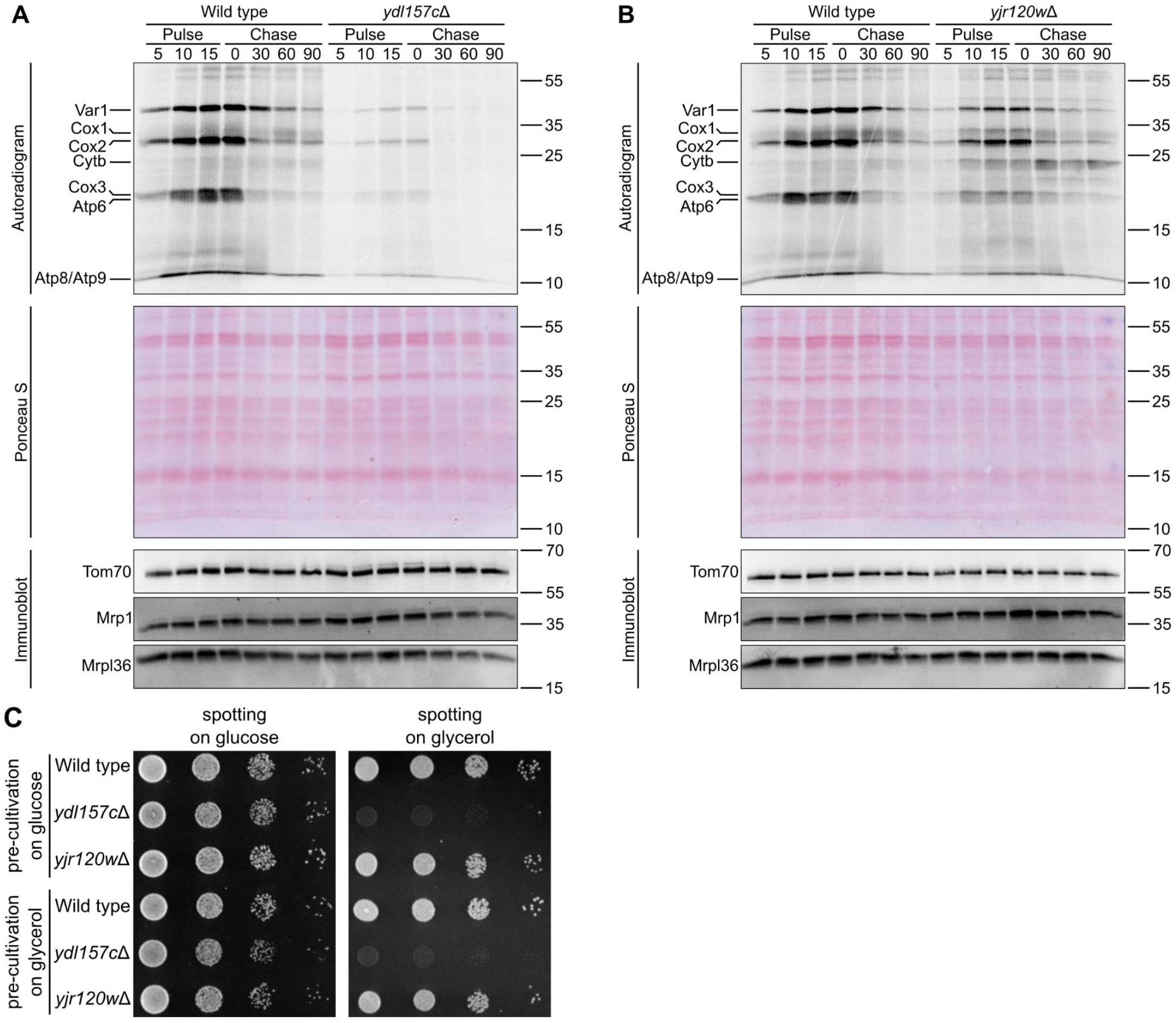
Additional Evidence Supporting Mitochondrial Translation MIMaL connections. Wild type and *ydl157cΔ* **(A)** or *yjr120wΔ* **(B)** cells were treated with cycloheximide and mitochondrial translation products were labeled with 35S-methionine for 15 minutes (pulse) at 30°C. The labeling was stopped by adding excess cold methionine and the temperature was increased to 37°C to induce protein destabilization (chase for a total of 90 min). Loading controls are included. Differences in translation between the strains reflect the connection between *YJR120W* and the mitochondrial ribosome genes such as *MRP1*. **(C)** Each strain was tested for their ability to grow under respiration conditions by comparing spotting on YPD and YPG.

**Supplemental Figure 4.**
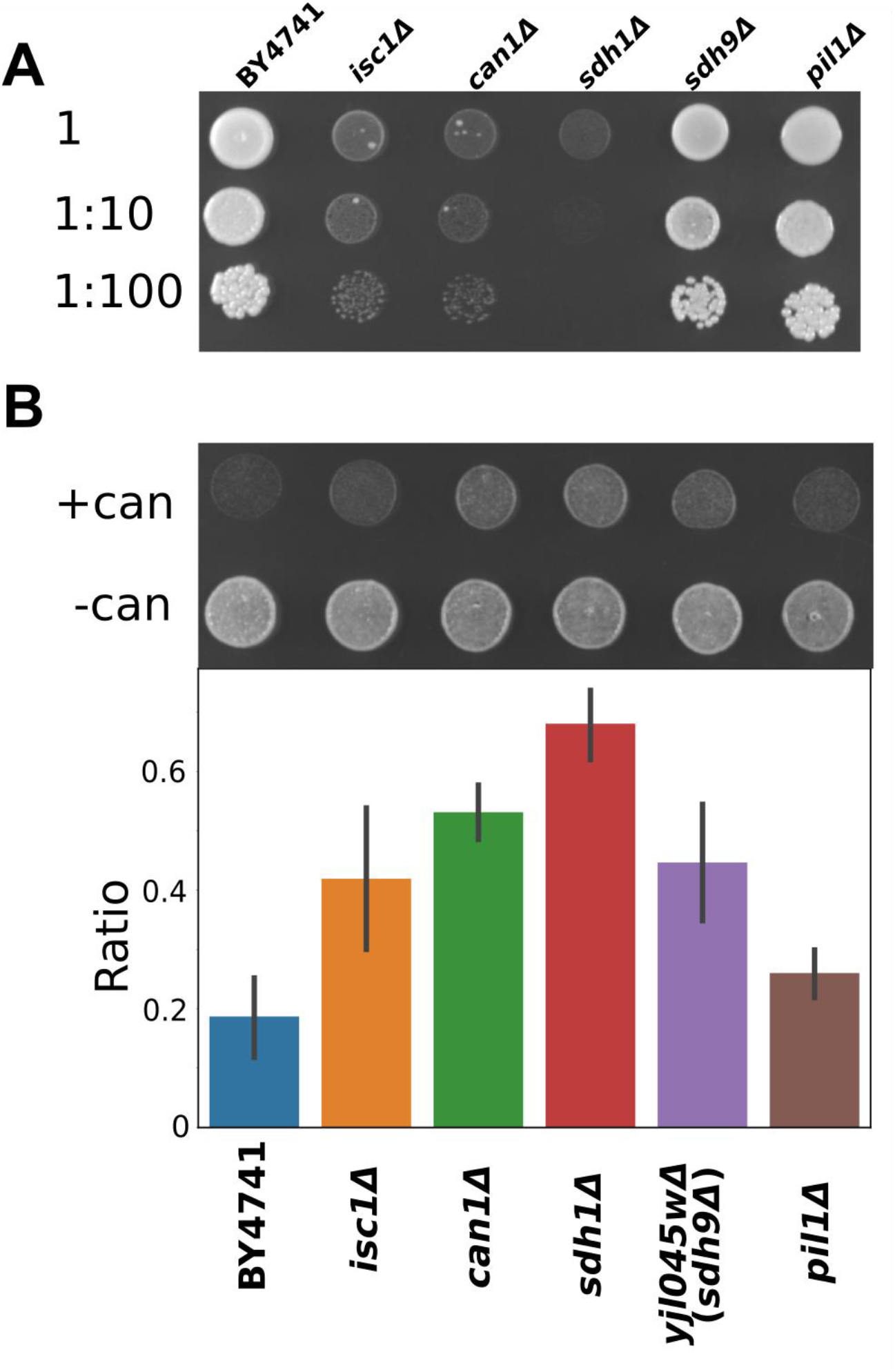
Additional Evidence Supporting Eisosomal MIMaL connections. **(A)** Growth of strains related to *pil1Δ* on SC -arg. **(B)** Resistance to canavanine was tested in strains related to *pil1Δ* by exposure to high concentrations of canavanine over 72 hours. Growth of each strain was tested before and after exposure and quantified using imagej. Significant differences (p-value <= 0.024, Tukey’s HSD) in growth were seen between wild-type and all other strains, except *pil1Δ*.

**Supplemental Figure 5.**
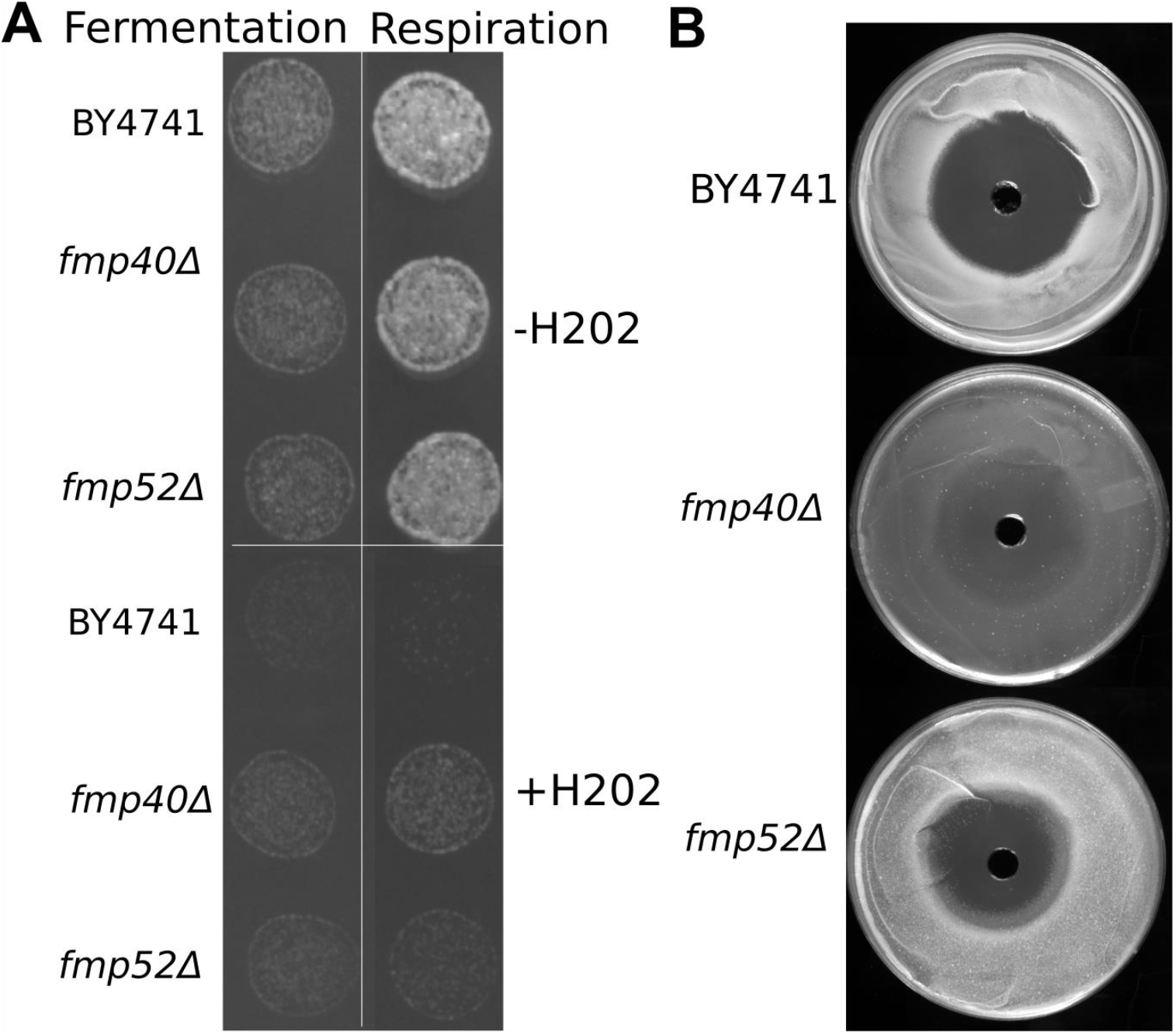
Additional Evidence Supporting Oxidative Stress Response MIMaL connections. **(A)** The strains *fmp40Δ* and *fmp52Δ* were tested for resistance to hydrogen peroxide stress under fermentation and respiration conditions. Resistance was quantified by calculating the ratio of growth before hydrogen peroxide treatment to growth after hydrogen peroxide treatment. Growth was quantified by measuring the greyscale of a circular area from the center of each drop **(Supplemental Information). (B)** Resistance to hydrogen peroxide was assessed by a zone of inhibition assay on YPG for *fmp40Δ* and *fmp52Δ*. Growth closer to the source of hydrogen peroxide is seen in both *fmp40Δ* and *fmp52Δ* when compared to wild type.

**Supplemental Figure 6.**
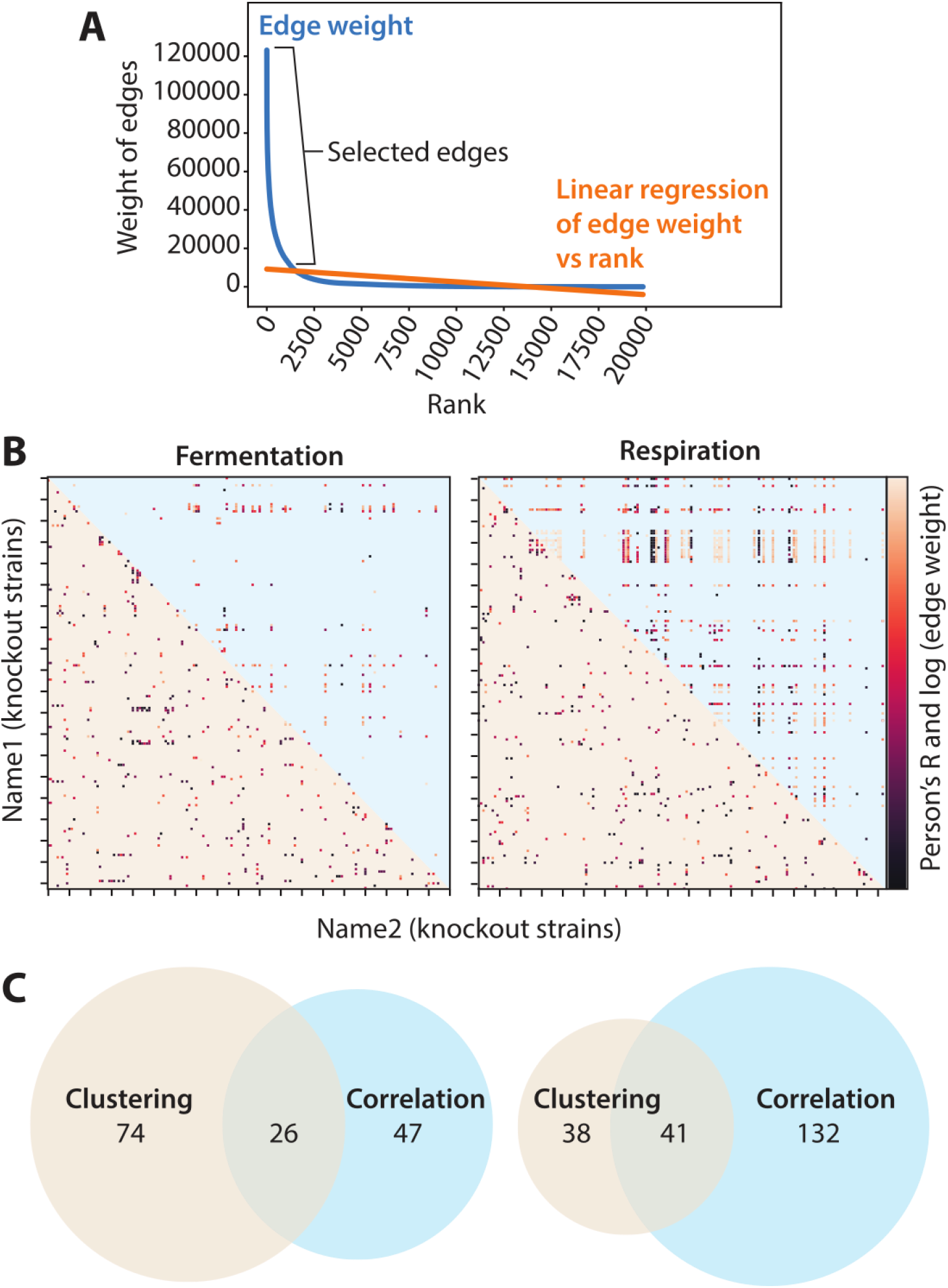
MIMaL Recapitulates Unique Set of Known Connections. **(A)** From the set of all connections between knockouts across all metabolites, the most important were selected by calculating a linear regression between the weight and the rank of the value. All connections with a weight above 8210 were kept for further analysis. **(B)** Weights calculated from MIMAL were compared with proteome-proteome correlations from the Y3K dataset for both respiration and fermentation. **(C)** Connections between knockouts were compared with correlations between knockouts in identifying known relationships between genes. A set of all known positive and negative genetic interactions and physical interactions was compared with the set of connections and correlations. A unique set of interactions was recapitulated from both MIMaL and correlations.

