## supplemental tables for "Multi-Omic Integration by Machine Learning (MIMaL) Reveals Protein-Metabolite Connections and New Gene Functions": supplemental tables.docx

**Supplementary tables**

Supplementary table 4: Yeast strains used in this study.

| **Strain** | **Genotype** | **Source** |
| --- | --- | --- |
| W303 wild type | *MAT a {leu2-3,112 trp1-1 can1-100 ura3-1 ade2-1 his3-11,15}* | Euroscarf |
| W303 *yjr120w*Δ | *MAT a {leu2-3,112 trp1-1 can1-100 ura3-1 ade2-1 his3-11,15} YJR120W::HIS3MX6* | This study |
| W303 ydl*157c*Δ | *MAT a {leu2-3,112 trp1-1 can1-100 ura3-1 ade2-1 his3-11,15} YDL157C::HIS3MX6* | This study |
| BY4743 | *MATa his3Δ1 leu2Δ0 met15Δ0 ura3Δ0/MATalpha his3Δ1 leu2Δ0 lys2Δ0 ura3Δ0* | Horizon Discovery |
| BY4743 AAT2Δ | *MATa his3Δ1 leu2Δ0 met15Δ0 ura3Δ0 AAT2Δ /MATalpha his3Δ1 leu2Δ0 lys2Δ0 ura3Δ0 AAT2Δ* | Horizon Discovery |
| BY4743 ALD5Δ | *MATa his3Δ1 leu2Δ0 met15Δ0 ura3Δ0 ALD5Δ /MATalpha his3Δ1 leu2Δ0 lys2Δ0 ura3Δ0 ALD5Δ* | Horizon Discovery |
| BY4743 MEF1Δ | *MATa his3Δ1 leu2Δ0 met15Δ0 ura3Δ0 MEF1Δ /MATalpha his3Δ1 leu2Δ0 lys2Δ0 ura3Δ0 MEF1Δ* | Horizon Discovery |
| BY4741 | *MATa his3Δ1 leu2Δ0 met15Δ0 ura3Δ0* | Horizon Discovery |
| BY4741 CAN1Δ | *MATa his3Δ1 leu2Δ0 met15Δ0 ura3Δ0 CAN1Δ* | Horizon Discovery |
| BY4741 ISC1Δ | *MATa his3Δ1 leu2Δ0 met15Δ0 ura3Δ0 ISC1Δ* | Horizon Discovery |
| BY4741 SDH1Δ | *MATa his3Δ1 leu2Δ0 met15Δ0 ura3Δ0 SDH1Δ* | Horizon Discovery |
| BY4741 SDH9Δ | *MATa his3Δ1 leu2Δ0 met15Δ0 ura3Δ0 SDH9Δ* | Horizon Discovery |
| BY4741 PIL1Δ | *MATa his3Δ1 leu2Δ0 met15Δ0 ura3Δ0 PIL1Δ* | Horizon Discovery |
| BY4741 FMP40Δ | *MATa his3Δ1 leu2Δ0 met15Δ0 ura3Δ0 FMP40Δ* | Horizon Discovery |
| BY4741 FMP52Δ | *MATa his3Δ1 leu2Δ0 met15Δ0 ura3Δ0 FMP52*Δ | Horizon Discovery |
| BY4741 SET4Δ | *MATa his3Δ1 leu2Δ0 met15Δ0 ura3Δ0 SET4Δ* | Horizon Discovery |

Supplementary table 5: Oligonucleotides and PCR templates used for chromosomal modification and respective control experiments.

| **Modification** | **Oligonucleotides** | **PCR template** |
| --- | --- | --- |
| Deletion of *YJR120W* | 5’- AATACTCTAAACCAGTAATAAAGCCACATATATATGTATTTCCTTTCACGTGATGCGTACGCTGCAGGTCGAC-3‘  5‘- AGACAAATTTGACTTGAGGAGGAGCACTTTTGTCTTCGTTTATGAGGGGGGTTTAATCGATGAATTCGAGCTCG -3‘ | *pFA6a–HisMX6* |
| Control PCR for *yjr1202*Δ | 5‘- GTCGACCTGCAGCGTACG -3‘  5‘- CGGCTGTTTCTGTGAGCGG -3‘ | Colony PCR of transformants |
| Deletion of *YDL157C* | 5’- CAAGTAGGAAACAAAACACCGCTTAGAGAGAAACAGCAAGTGGTGAAAAACAATGCGTACGCTGCAGGTCGAC -3‘  5‘- TAGCACTTTATGTACAGAAAATTTCACGATTGAGAGAAGCTTGCGAATAATACTAATCGATGAATTCGAGCTCG -3‘ | *pFA6a–HisMX6* |
| Control PCR for *ydl157c*Δ | 5‘- GTCGACCTGCAGCGTACG -3‘  5’- CACATCGTCTGCAGACTCACC -3‘ | Colony PCR of transformants |
